# A haplotype-resolved, *de novo* genome assembly for the wood tiger moth (*Arctia plantaginis*) through trio binning

**DOI:** 10.1101/2020.02.28.970020

**Authors:** Eugenie C. Yen, Shane A. McCarthy, Juan A. Galarza, Tomas N. Generalovic, Sarah Pelan, Petr Nguyen, Joana I. Meier, Ian A. Warren, Johanna Mappes, Richard Durbin, Chris D. Jiggins

## Abstract

**Background:** Diploid genome assembly is typically impeded by heterozygosity, as it introduces errors when haplotypes are collapsed into a consensus sequence. Trio binning offers an innovative solution which exploits heterozygosity for assembly. Short, parental reads are used to assign parental origin to long reads from their F1 offspring before assembly, enabling complete haplotype resolution. Trio binning could therefore provide an effective strategy for assembling highly heterozygous genomes which are traditionally problematic, such as insect genomes. This includes the wood tiger moth (*Arctia plantaginis*), which is an evolutionary study system for warning colour polymorphism.

**Findings:** We produced a high-quality, haplotype-resolved assembly for *Arctia plantaginis* through trio binning. We sequenced a same-species family (F1 heterozygosity ∼1.9%) and used parental Illumina reads to bin 99.98% of offspring Pacific Biosciences reads by parental origin, before assembling each haplotype separately and scaffolding with 10X linked-reads. Both assemblies are highly contiguous (mean scaffold N50: 8.2Mb) and complete (mean BUSCO completeness: 97.3%), with complete annotations and 31 chromosomes identified through karyotyping. We employed the assembly to analyse genome-wide population structure and relationships between 40 wild resequenced individuals from five populations across Europe, revealing the Georgian population as the most genetically differentiated with the lowest genetic diversity.

**Conclusions:** We present the first invertebrate genome to be assembled via trio binning. This assembly is one of the highest quality genomes available for Lepidoptera, supporting trio binning as a potent strategy for assembling highly heterozygous genomes. Using this assembly, we provide genomic insights into geographic population structure of *Arctia plantaginis.*

## DATA DESCRIPTION

### Background

The ongoing explosion in *de novo* reference genome assembly for non-model organisms has been facilitated by the combination of advancing technologies and falling costs of next generation sequencing [1]. Long-read sequencing technologies further revolutionised the quality of assembly achievable, with incorporation of long reads that can span common repetitive regions leading to radical improvements in contiguity [2]. However, heterozygosity still presents a major challenge to *de novo* assembly of diploid genomes. Most current technologies attempt to collapse parental haplotypes into a composite, haploid sequence, introducing erroneous duplications through mis-assembly of heterozygous sites as separate genomic regions. This problem is exacerbated in highly heterozygous genomes, resulting in fragmented and inflated assemblies which impede downstream analyses [3, 4]. Furthermore, a consensus sequence does not represent either true, parental haplotype, leading to loss of haplotype-specific information such as allelic and structural variants [5]. Whilst reducing heterozygosity by inbreeding has been a frequent approach, rearing inbred lines is unfeasible and highly time consuming for many non-model systems, and resulting genomes may no longer be representative of wild populations.

Trio binning is an innovative, new approach which takes advantage of heterozygosity instead of trying to remove it [6]. In this method, a family trio is sequenced with short reads for both parents and long reads for an F1 offspring. Parent-specific k-mer markers are then identified from the parental reads and used to assign offspring reads into maternal and paternal bins, before assembling each parental haploid genome separately [6]. The ability of trio binning to accurately distinguish parental haplotypes increases at greater heterozygosity, with high-quality, *de novo* assemblies achieved for bovid genomes by crossing different breeds [6] and species [7] to maximise heterozygosity. Therefore, trio binning has the potential to overcome current difficulties faced by highly heterozygous genomes, which have typically evaded high-quality assembly through conventional methods.

We utilised trio binning to assemble a high-quality, haplotype-resolved reference genome for the wood tiger moth (*Arctia plantaginis*; formerly *Parasemia plantaginis* [8]). This represents the first trio binned assembly available for Insecta and indeed any invertebrate animal species, diversifying the organisms for which trio binning has been applied outside of bovids [6, 7], zebra finches [9], humans [6, 9, 10] and *Arabidopsis thaliana* [6]. Using a family trio with same-species *A. plantaginis* parents, 99.98% of offspring reads were successfully binned into parental haplotypes. This was possible due to the high heterozygosity of the *A. plantaginis* genome; heterozygosity of the F1 offspring was estimated to be ∼1.9%, exceeding levels (∼1.2%) obtained when crossing different bovid species [7]. Both resulting haploid assemblies are highly contiguous and complete, strongly supporting trio binning as an effective strategy for *de novo* assembly of heterozygous genomes.

The presented *A. plantaginis* assembly will also provide an important contribution to the growing collection of lepidopteran reference genomes [11]. Comparative phylogenomic studies will benefit from the addition of *A. plantaginis* to the phylogenomic dataset [12,13], being the first species to be sequenced within the Erebidae family [8, 14], and the first fully haplotype-resolved genome available for Lepidoptera. *A. plantaginis* itself is an important evolutionary study system, being a moth species which uses aposematic hindwing colouration to warn avian predators of its unpalatability [15]. Whilst female hindwing colouration varies continuously from orange to red, male hindwings exhibit a discrete colour polymorphism maintained within populations (Figure 1), varying in frequency from yellow-white in Europe and Siberia, yellow-red in the Caucasus, and black-white in North America and Northern Asia [16, 17]. Hence, *A. plantaginis* provides a natural system to study the evolutionary forces that promote phenotypic diversification on local and global scales, for which availability of a high-quality, haplotype-resolved and annotated reference genome will now transform genetic research.

**Figure 1.**
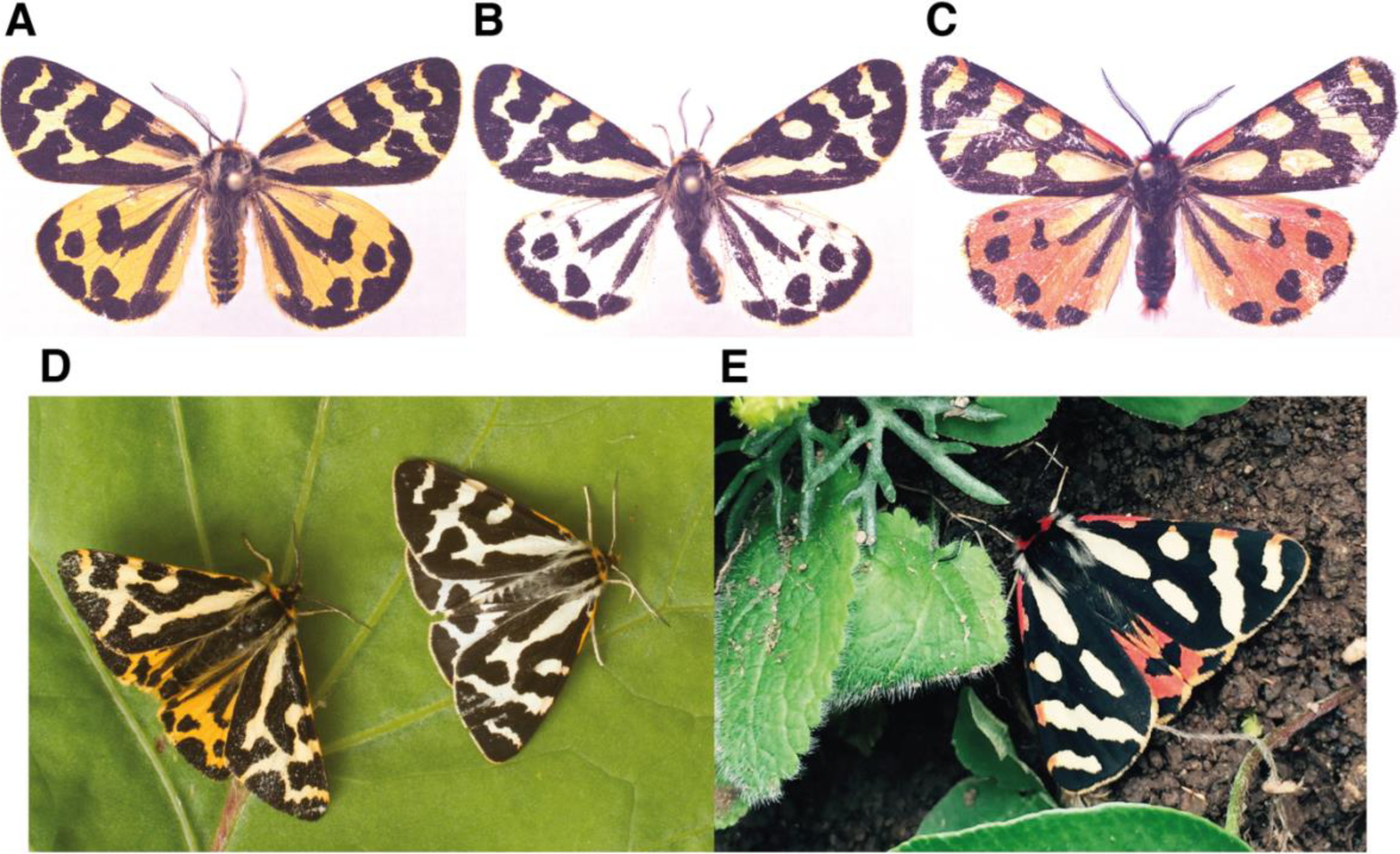
Discrete colour morphs of *Arctia plantaginis* males. Whilst forewings remain white, hindwings are polymorphic with variable black patterns, existing as discrete (**A)** yellow (**B)** white and (**C)** red morphs, which can only be found in the Caucasus region. **(A-C)** show pinned dead morphs **(D-E)** show examples of morphs in the wild. Photos: Johanna Mappes and Ossi Nokelainen.

## Materials and Methods

### Cross preparation and sequencing

To obtain an *A. plantaginis* family trio, selection lines for yellow and white male morphs were created from Finnish populations at the University of Jyväskylä over three consecutive generations. Larvae were fed with wild dandelion (Taraxacum spp.) and reared under natural light conditions, with an average temperature of 25°C during the day and 15-20°C at night until pupations. A father from the white selection line and mother from the yellow selection line were crossed, then collected and dry-frozen along with their F1 pupae at −20°C in 1.5 ml (millilitre) sterile Eppendorf tubes.

For short-read sequencing of the father (sample ID: CAM015099; ENA accession number: ERS4285278) and mother (sample ID: CAM015100; ENA accession number: ERS4285279), DNA was extracted from adult thoraces using a QIAGEN DNeasy Blood & Tissue Kit (Qiagen, Germany) following the manufacturer’s protocol, then library preparation and sequencing was performed by Novogene (China). Illumina NEBNext (New England Biolabs, United States) libraries were constructed with an insert size of 350 bp (base pair), following the manufacturer’s protocol, and sequenced with 150 bp paired end reads on an Illumina NovaSeq 6000 platform (Illumina, United States).

For long-read sequencing of a single F1 pupal offspring (Sample ID: CAM015101; ENA accession number: ERS4285595), high-molecular weight DNA was extracted from the entire body of one F1 pupa using a QIAGEN Blood & Culture DNA Midi Kit (Qiagen, Germany) following the manufacturer’s protocol, then library preparation and sequencing was performed by the Wellcome Sanger Institute (Cambridge, UK). A SMRTbell CLR (continuous long reads) sequencing library was constructed following the manufacturer’s protocol, and sequenced on 5 SMRT (Single Molecule Real-Time) cells within a PacBio Sequel system (Pacific Biosciences, United States) using version 3.0 chemistry and 10 hour runs. This generated 3,474,690 subreads, with a subread N50 of 18.8 kb and total of 39,471,717,610 bp. From the same sample, a 10X Genomics Chromium linked-read sequencing library (10X Genomics, United States) was also prepared following the manufacturer’s protocol, and sequenced with 150 bp paired end reads on an Illumina HiSeq X Ten platform (Illumina, United States). This generated 625,914,906 reads, and after mapping to the assembly described below, we estimate a barcoded molecule length of ∼43 kbp.

### Trio binning genome assembly

Canu version 1.8 [18] was used to bin *A. plantaginis* F1 offspring PacBio (Pacific Biosciences) subreads into those matching the paternal and maternal haplotypes defined by k-mers specific to the maternal and paternal Illumina data (Supplementary Figure 1). This resulted in 1,662,000 subreads assigned to the paternal haplotype, 1,529,779 subreads assigned to the maternal haplotype, and 2,445 (0.07%) subreads unassigned. Using only the assigned reads, the haplotype binned reads were assembled separately using wtdbg2 version 2.3 [19], with the ‘-xsq’ pre-set option for PacBio Sequel data and an estimated genome size of 550Mb. The assemblies were polished using Arrow version 2.3.3 [20] and the haplotype binned PacBio reads. The 10X linked-reads were then used to scaffold each assembly using scaff10x [21], followed by another round of Arrow polishing on the scaffolds. To polish further with the 10X linked-read Illumina data, we first concatenated the two scaffolded assemblies, mapped the 10X Illumina data with Long Ranger version 2.2.0 [22] longranger align, called variants with freebayes version 1.3.1 [23], then applied homozygous non-reference edits to the assembly using bcftools consensus [24]. The assembly was then split back into paternal and maternal components, giving separate paternal haplotype (iArcPla.TrioW) and maternal haplotype (iArcPla.TrioY) assemblies.

The assemblies were checked for contamination and further manually assessed and corrected using gEVAL [25]. The Kmer Analysis Toolkit (KAT) version 2.4.2 [26] was used to compare k-mers from the 10X Illumina data to k-mers in each of the haplotype-resolved assemblies, and in the combined diploid assembly representing both haplotypes. Phasing of the assembled contigs and scaffolds was visualised using the parental k-mer databases produced by Canu [27]. Haploid genome size, heterozygosity and repeat fraction of the F1 offspring were estimated using GenomeScope [28] and k-mers derived from the 10X Illumina data.

### Quality assessment

To assess the quality of each parental haplotype of the *A. plantaginis* trio binned assembly, standard contiguity metrics were computed, and assembly completeness was evaluated by calculating BUSCO (Benchmarking Universal Single-Copy Ortholog) scores using BUSCO version 3.0.2, comparing against the ‘insecta_odb9’ database of 1658 Insecta BUSCO genes with default Augustus parameters [29]. Quality comparisons were conducted against an assembly of unbinned data from the same F1 offspring (iArcPla.wtdbg2), and against a representative selection of published lepidopteran reference genomes. For this, the latest versions of seven Lepidoptera species were downloaded: *Bicyclus anynana* version 1.2 [30], *Danaus plexippus* version 3 [31], *Heliconius melpomene* version Hmel.2.5 [32], *Manduca sexta* version Msex_1.0 [33] and *Melitaea cinxia* version MelCinx1.0 [34] were downloaded from Lepbase version 4.0 [11], whilst *Bombyx mori* version Bomo_genome_assembly [35] was downloaded from SilkBase version 2.1 [36] and *Trichoplusia ni* version PPHH01.1 [37] was downloaded from NCBI RefSeq version 94 [38]. Cumulative scaffold plots were visualised in R version 3.5.1 [39] using the ggplot2 package version 3.1.1 [40].

### Genome annotation

Genome annotations were produced for each parental haplotype of the *A. plantaginis* trio binned assembly using the BRAKER2 version 2.1.3 pipeline [41]. A *de novo* library of repetitive sequences was identified with both genomes using RepeatScout version 1.0.5 [42]. Repetitive regions of the genomes were soft masked using RepeatMasker version 4.0.9 [43], Tandem Repeats Finder version 4.00 [44] and the RMBlast version 2.6.0 sequence search engine [45] combined with the Dfam_Consensus-20170127 database [46]. Raw RNA-seq reads were obtained from Galarza et al. 2017 [47] under study accession number PRJEB14172, and arthropod proteins were obtained from OrthoDB [48]. RNA-seq reads were trimmed for adapter contamination using cutadapt version 1.8.1 [49] and quality controlled pre and post trimming with fastqc version 0.11.8 [50]. RNA-seq reads were mapped to each respective genome using STAR (Spliced Transcripts Alignment to a Reference) version 2.7.1 [51]. Arthropod proteins were aligned to the genomes using GenomeThreader version 1.7.0 [52]. BRAKER2’s *ab initio* gene predictions were carried out using homologous protein and *de novo* RNA-seq evidence using Augustus version 3.3.2 [41] and GeneMark-ET version 4.38 [41]. Annotation completeness was assessed using BUSCO version 3.0.2 against the ‘insecta_odb9’ database of 1658 Insecta BUSCO genes with default Augustus parameters [29].

### Cytogenetic analysis

Spread chromosome preparations for cytogenetic analysis were produced from wing imaginal discs and gonads of third to fifth instar larvae, according to Šíchová et al. 2013 [53]. Female and male gDNA were extracted using the CTAB (hexadecyltrimethylammonium bromide) method, adapted from Winnepenninckx et al. 1993 [54]. These were used to generate probe and competitor DNA, respectively, for genomic *in situ* hybridization (GISH). Female genomic probe was labelled with Cy3-dUTP (cyanine 3-deoxyuridine triphosphate; Jena Bioscience, Germany) by nick translation, following Kato et al. 2006 [55] with a 3.5 hour incubation at 15°C. Male competitor DNA was fragmented with a 20 minute boil. GISH was performed following the protocol of Yoshido et al. 2005 [56]. For each slide, the hybridization cocktail contained 250 ng of female labelled probe, 2-3 µg of male competitor DNA, and 25 µg of salmon sperm DNA. Preparations were counterstained with 0.5 mg/ml DAPI (4’,6-diamidino-2-phenylindole; Sigma-Aldrich) in DABCO antifade (1,4-diazabicyclo[2.2.2]octane; Sigma-Aldrich). Results were observed in the Zeiss Axioplan 2 Microscope (Carl Zeiss, Germany) and documented with an Olympus CCDMonochrome Camera XM10, with the cellSens 1.9 digital imaging software (Olympus Europa Holding, Germany). Images were pseudo-colored and superimposed in Adobe Photoshop CS3.

### Population genomic analysis

We implemented the novel *A. plantaginis* reference assembly to analyse patterns of population genomic variation between 40 wild, adult males sampled from the European portion of *A. plantaginis’* Holarctic species range [17]. Samples were collected by netting and pheromone traps from Central Finnish (n=10) and Southern Finnish populations (n=10) where yellow and white morphs exist in equal proportions, an Estonian population (n=5) where white morphs are frequent compared to rare yellow morphs, a Scottish population (n=10) where only yellow morphs exist, and a Georgian population (n=5) where red morphs exist alongside yellow morphs (Figure 5A). Exact sampling localities are available in Supplementary Table 2. DNA was extracted from thoraces using a QIAGEN DNeasy Blood & Tissue Kit (Qiagen, Germany) following the manufacturer’s protocol, then library preparation and sequencing was performed by Novogene (China). Illumina NEBNext (New England Biolabs, United States) libraries were constructed with an insert size of 350 bp, following the manufacturer’s protocol, and sequenced with 150 bp, paired end reads on an Illumina NovaSeq 6000 platform (Illumina, United States). ENA accession numbers for all resequenced samples are available in Supplementary Table 3.

Reads were mapped against the paternal iArcPla.TrioW assembly (chosen due to higher assembly completeness; Table 2) using BWA-MEM (Burrows-Wheeler Aligner) version 7.17 [57] with default parameters, resulting in a mean sequencing coverage of 13X (Supplementary Table 3). Alignments were sorted with SAMtools version 1.9 [58] and PCR-duplicates were removed with Picard version 2.18.15 [59]. Variants were called for each sample using Genome Analysis Tool Kit (GATK) HaplotypeCaller version 3.7 [60, 61], followed by joint genotyping across all samples using GATK version 4.1 GenotypeGVCFs [60, 61], with expected heterozygosity set to 0.01. The raw SNP (single nucleotide polymorphism) callset was quality filtered by applying thresholds: quality by depth (QD>2.0), root mean square mapping quality (MQ>50.0), mapping quality rank sum test (MQRankSum>-12.5), read position rank sum test (ReadPosRankSum>-8.0), Fisher strand bias (FS<60.0) and strand odds ratio (SOR<3.0). Filters by depth (DP) of greater than half the mean (DP>409X) and less than double the mean (DP<1636X) were also applied. Linkage disequilibrium (LD) pruning was applied using the ldPruning.sh script [62] with an LD threshold of r^2^<0.01, in 50kb windows shifting by 10kb. This callset was further filtered for probability of heterozygosity excess p-value>1×10^−5^ using VCFtools version 0.1.15 [63] to exclude potential paralogous regions, giving an analysis-ready callset.

**Table 1.** Genome annotation statistics for the *Arctia plantaginis* trio binned assembly. Statistics generated using the BRAKER2 pipeline, for the paternal (iArcPla.TrioW) and maternal (iArcPla.TrioY) haplotype assemblies.

|  |  | iArcPla.TrioW (paternal) | iArcPla.TrioY (maternal) |
| --- | --- | --- | --- |
| Total Genome size (bp) |  | 584,621,344 | 577,993,050 |
| Repetitive sequences (bp) |  | 239,949,688 | 247,356,128 |
| Masked repeats (%) |  | 41.04 | 42.80 |
| Mapped RNA-seq reads (n) |  | 599,065,138 | 590,780,528 |
| Mapped RNA-seq reads (%) |  | 95.45 | 94.13 |
| Protein-coding genes (n) |  | 19,899 | 18,894 |
| Mean gene length (bp) |  | 5,966 | 5,951 |
| BUSCO Completeness (%; n:1658) |  | 98.00 | 95.90 |
| Repeat Elements (n) | Total | 11,320 | 12,576 |
|  | DNA Transposons | 3,222 | 3,366 |
|  | LTR | 1,891 | 2,192 |
|  | LINES | 3,006 | 3,506 |
|  | SINES | 544 | 547 |
|  | Unclassified | 2,657 | 2,965 |

**Table 2.** Comparison of assembly contiguity and completeness between *Arctia plantaginis* and seven publicly available lepidopteran assemblies. Standard contiguity and BUSCO completeness metrics generated for each genome assembly, highlighting the high-quality *A. plantaginis* assembly achieved by trio binning. See **Figure 3** for assembly contiguity visualisation via cumulative scaffold plots, and **Supplementary Table 1** for the full BUSCO analysis summary.

|  | Assembly contiguity |  |  |  |  | Assembly completeness |  |  |
| --- | --- | --- | --- | --- | --- | --- | --- | --- |
|  | Assembly size (Mb) | Total scaffolds | Longest scaffold (Mb) | N50 (kb) | N50 count | Total complete BUSCOs | Single copy BUSCOs | Duplicated BUSCOs |
| *Arctia plantaginis*<br>(binned: iArcPla.TrioW, paternal haplotype) | 585 | 1069 | 21.5 | 6730 | 24 | 98.1% | 96.9% | 1.2% |
| *Arctia plantaginis*<br>(binned: iArcPla.TrioY, maternal haplotype) | 578 | 1050 | 24.4 | 9770 | 18 | 96.4% | 95.3% | 1.1% |
| *Arctia plantaginis*<br>(unbinned: iArcPla.wtdbg2) | 615 | 2948 | 11.3 | 1840 | 85 | 95.4% | 93.3% | 2.1% |
| *Bicyclus anynana* | 475 | 10800 | 5.04 | 638.3 | 194 | 97.6% | 96.8% | 0.8% |
| *Bombyx mori* | 482 | 696 | 21.5 | 16796 | 13 | 98.4% | 97.2% | 1.2% |
| *Danaus plexippus* | 249 | 5397 | 6.24 | 715.6 | 101 | 98.0% | 96.0% | 2.0% |
| *Heliconius melpomene* | 275 | 332 | 18.1 | 14308 | 9 | 98.7% | 97.4% | 1.3% |
| *Manduca sexta* | 419 | 20871 | 3.25 | 664.0 | 169 | 96.7% | 93.9% | 2.8% |
| *Melitaea cinxia* | 390 | 8261 | 0.668 | 119.3 | 970 | 83.0% | 82.9% | 0.1% |
| *Trichoplusia ni* | 333 | 1916 | 8.93 | 4648 | 27 | 97.4% | 96.6% | 0.8% |

An unrooted, maximum likelihood (ML) phylogenetic tree was constructed to evaluate phylogenomic relationships, using the analysis-ready callset which was further reduced in size by subsampling every other SNP. The best-scoring ML tree was built in RAxML (Random Axelerated Maximum Likelihood) version 8.2.12 [64] with 100 rapid bootstrap replicates, using the GTRGAMMA model (generalised time-reversible substitution model and gamma model of rate heterogeneity) and Lewis ascertainment bias correction to account for the lack of monomorphic sites, then visualised in FigTree version 1.4.4 [65]. A principle component analysis (PCA) was also conducted to evaluate genome-wide population structure. A minor allele frequency filter of 0.05 was applied to the analysis-ready callset using VCFtools version 0.1.15 [63] to remove PCA-uninformative SNPs, then PCA was performed in R version 3.5.1 [39] using the SNPRelate package version 3.3 [66].

## Results and Discussion

### Trio binning genome assembly

K-mer spectra plots (Figure 2) indicate a highly complete assembly of both parental haplotypes in the *A. plantaginis* diploid offspring genome. There is good separation between the parental haplotypes, as each haploid assembly consists mostly of single-copy k-mers with low frequency of 2-copy k-mers, indicating a correctly haplotype-resolved assembly with low levels of artefactual duplication (Figure 2B, 2C; Supplementary Figure 2). This is also confirmed by the spectra plot for the combined diploid assembly (Figure 2A), where homozygous regions consist mostly of 2-copy k-mers and heterozygous regions consist mostly of 1-copy k-mers, as expected from the presence of both complete, parental haplotypes and low artefactual duplication. Using GenomeScope, we estimated the F1 offspring haploid genome size to be 590Mb with a repeat fraction of 27% (Supplementary Figure 3). Successful haplotype separation was possible due to the high estimated heterozygosity (∼1.9%) of the F1 offspring genome (Supplementary Figure 3), with greater levels of heterozygosity achieved through our same-species *A. plantaginis* cross than previously achieved through an inter-species cross between yak (*Bos grunniens*) and cattle *(Bos taurus*), which gave an F1 heterozygosity of ∼1.2% [7].

**Figure 2.**
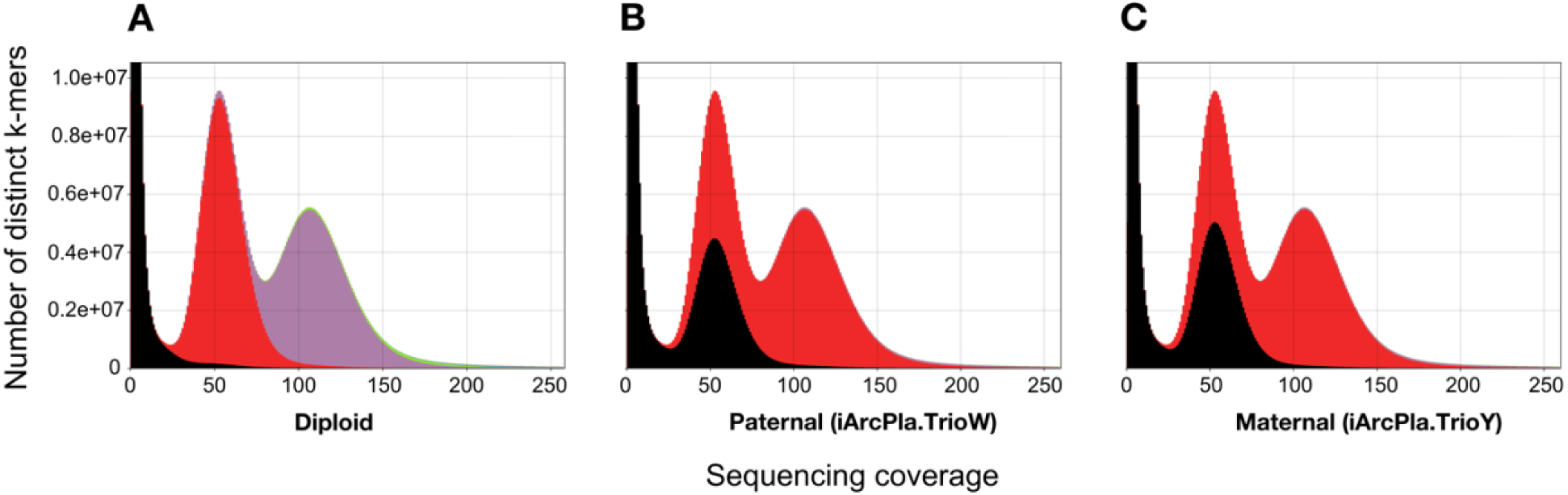
K-mer spectra plots for the *Arctia plantaginis* trio binned genome assembly. Plots produced using K-mer Analysis Toolkit (KAT), showing the frequency of k-mers in an assembly versus the frequency of k-mers (i.e. sequencing coverage) in the 10X Illumina reads, for the **(A)** combined diploid assembly (paternal plus maternal), **(B)** paternal-only assembly (iArcPla.TrioW), and **(C)** maternal-only assembly (iArcPla.TrioY). Colours represent k-mer copy number in the assembly: black k-mers are not represented (0-copy), red k-mers are represented once (1-copy), purple k-mers are represented twice (2-copy) and green k-mers are represented thrice (3-copy). The first peak corresponds to k-mers missing from the assembly due to sequencing errors, the second peak corresponds to k-mers from heterozygous regions, and the third peak corresponds to k-mers from homozygous regions. These plots show a complete and well-separated assembly of both haplotypes in the F1 offspring diploid genome.

### Genome annotation

We identified and masked 222,866,714 bp (41.04%) and 227,797,418 bp (42.80%) of repetitive regions in the iArcPla.TrioW and iArcPla.TrioY assemblies, respectively (Table 1). The BRAKER2 pipeline annotated a total of 19,899 protein coding genes in the soft-masked iArcPla.TrioW genome with 98.0% BUSCO completeness, whilst 18,894 protein coding genes were annotated in the soft-masked iArcPla.TrioY genome with 95.9% BUSCO completeness (Table 1).

### Quality assessment

The paternal (iArcPla.TrioW) assembly contains 1069 scaffolds and N50=6.73 Mb, and the maternal (iArcPla.TrioY) assembly contains 1050 scaffolds and N50=9.77 Mb (Table 2). Both trio binned assemblies are more contiguous than the composite haploid iArcPla.wtdbg2 assembly produced using unbinned data from the same individual, which contains 2948 scaffolds and N50=1.84 Mb (Table 2; Figure 3A), illustrating the contiguity improvement we achieved by separating haplotypes before assembly. The trio binned assemblies are more complete than the unbinned assembly (complete BUSCOs: iArcPla.TrioW=98.1%; iArcPla.TrioY=96.4%; iArcPla.wtdbg2=95.4%). The trio binned assemblies are also less inflated than the unbinned assembly (assembly size: iArcPla.TrioW=585 Mb; iArcPla.TrioY=578 Mb; iArcPla.wtdbg2=615 Mb) and duplicated BUSCOs halved (duplicated BUSCOs: iArcPla.TrioW=1.2%; iArcPla.TrioY=1.1%; iArcPla.wtdbg2=2.1%), suggesting a reduction in artefactual assembly duplication at heterozygous sites through read binning (Table 2; Figure 3A).

**Figure 3.**
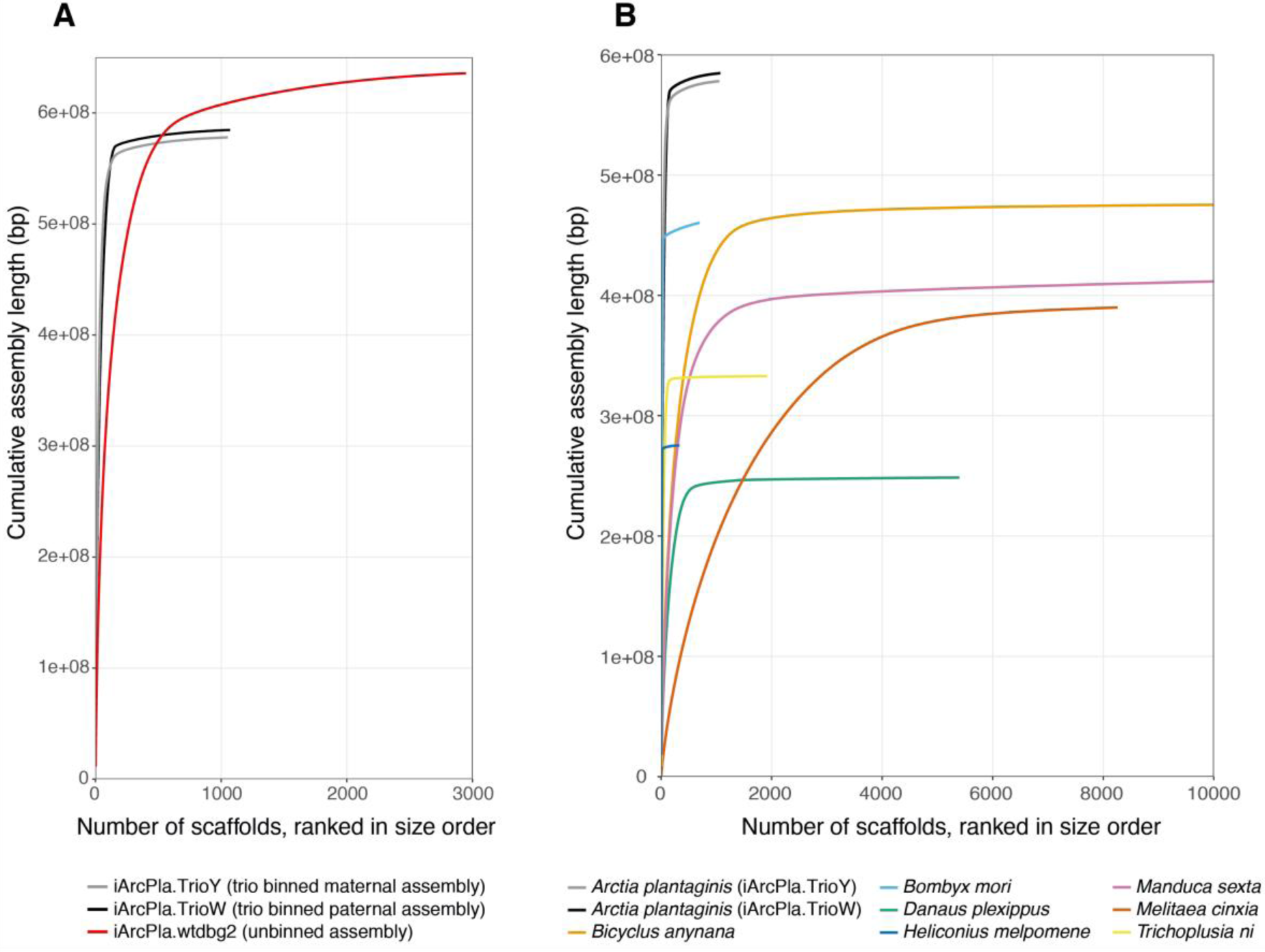
Cumulative scaffold plots visualise the high assembly contiguity of the trio binned *Arctia plantaginis* genome. A highly contiguous assembly is represented by a near vertical line with a short horizontal tail of trailing tiny scaffolds. **(A)** Comparison of the *A. plantaginis* trio binned assemblies iArcPla.TrioW (paternal haplotype) and iArcPla.TrioY (maternal haplotype) against the composite assembly using unbinned data from the same individual (iArcPla.wtdbg2). The much steeper curve and shorter horizontal tail for the trio binned assemblies compared to the unbinned assembly shows that trio binning greatly improved contiguity. **(B)** Comparison of the *A. plantaginis* trio binned assemblies against a representative selection of published lepidopteran genomes, shown up to the first 10000 scaffolds. This comparison demonstrates that the *A. plantaginis* trio binned assemblies are much more contiguous than most other lepidopteran genomes currently available.

The trio binned *A. plantaginis* assemblies are of comparable quality to the best reference genomes available for Lepidoptera (Table 2; Figure 3B). When compared to other published lepidopteran reference genomes, quality of the *A. plantaginis* assemblies surpasses all but the best *Heliconius melpomene* [32] and *Bombyx mori* [35] assemblies (Table 2; Figure 3B). As contiguity of the *H. melpomene* assembly was improved through pedigree linkage mapping and haplotypic sequence merging [32], whilst bacterial artificial chromosome (BAC) and fosmid clones were used to close gaps in the *B. mori* assembly [35], it is impressive that trio binning has instantly propelled contiguity of the *A. plantaginis* genome to very near that of *H. melpomene* and *B. mori*, before incorporating information from any additional technologies. Therefore, these comparisons strongly support trio binning as an effective strategy for *de novo* assembly of highly heterozygous genomes. Future chromosomal-level scaffolding work through Hi-C scaffolding technology [67] will elevate the *A. plantaginis* assembly quality to the top tier.

### Cytogenetic analysis

Mitotic nuclei prepared from wing imaginal discs of *A. plantaginis* larvae contained 2n=62 chromosomes in both sexes (Figure 4) in agreement with a previously reported modal chromosome number of arctiid moths [68], which is also the likely ancestral lepidopteran karyotype [34]. These insights will be helpful for future scaffolding work into a chromosomal-scale *A. plantaginis* reference assembly. Chromosomes decreased gradually in size, as is typical for lepidopteran karyotypes [69]. Due to the holokinetic nature of lepidopteran chromosomes, separation of sister chromatids by parallel disjunction was observed in mitotic metaphases [70]. Notably, two smallest chromosomes separated earlier compared to the other chromosomes (Figure 4A), although this could be an artefact of the spreading technique used for chromosome preparation. The presence of a W chromosome was confirmed in female nuclei by genomic *in situ* hybridization (Supplementary Figure 4; Supplementary Text 1).

**Figure 4.**
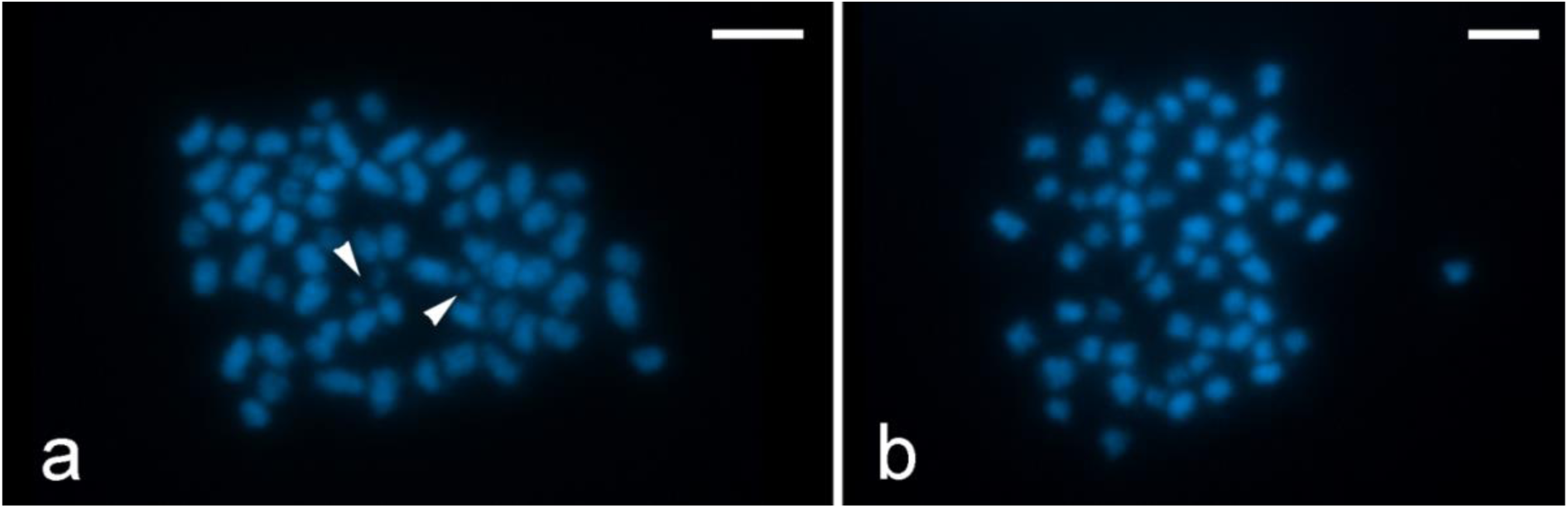
Cytogenetic analysis reveals 31 chromosomes in the *Arctia plantaginis* haploid genome. Chromosomes were counterstained with DAPI (blue). **(A)** Male mitotic metaphase consisted of 2n=62 chromosomes. Note separated chromatids of the smallest chromosome pair (arrowheads). **(B)** Female mitotic complement consisted of 2n=62 elements. Scale bar=5 µm.

### Population genomic variation across the European range

As an empirical application of the *A. plantaginis* reference genome, we conducted a population resequencing analysis to describe genomic variation between 40 wild *A. plantaginis* males from five populations spread across Europe (Figure 5A). PCA revealed clear population structuring with individuals clustering geographically by country of origin (Figure 5B), in congruence with strongly supported phylogenomic groupings also by country of origin (Figure 6). Central and Southern Finnish individuals grouped into a single population as expected from their geographic proximity (Figure B; Figure 6). The Finnish and Estonian populations clustered together away from the Scottish population along principle component (PC) 2 (Figure 5B) and on the phylogenetic tree (Figure 6), as would be predicted by effects of isolation by distance [71]. The Georgian population was highly genetically differentiated from all other sampled European populations, separating far along PC1 (Figure 5B) and possessing a much longer inter-population branch in the ML tree (Figure 6). Since the Georgian population has a distinctive genomic composition from the rest of the sampled distribution, this could support the hypothesis of incipient speciation in the Caucasus [17]. However, populations must be sampled in the large geographic gap between Georgia and the other populations in this preliminary analysis, to determine if genetic differentiation still persists when compared to nearby Central European populations.

**Figure 5.**
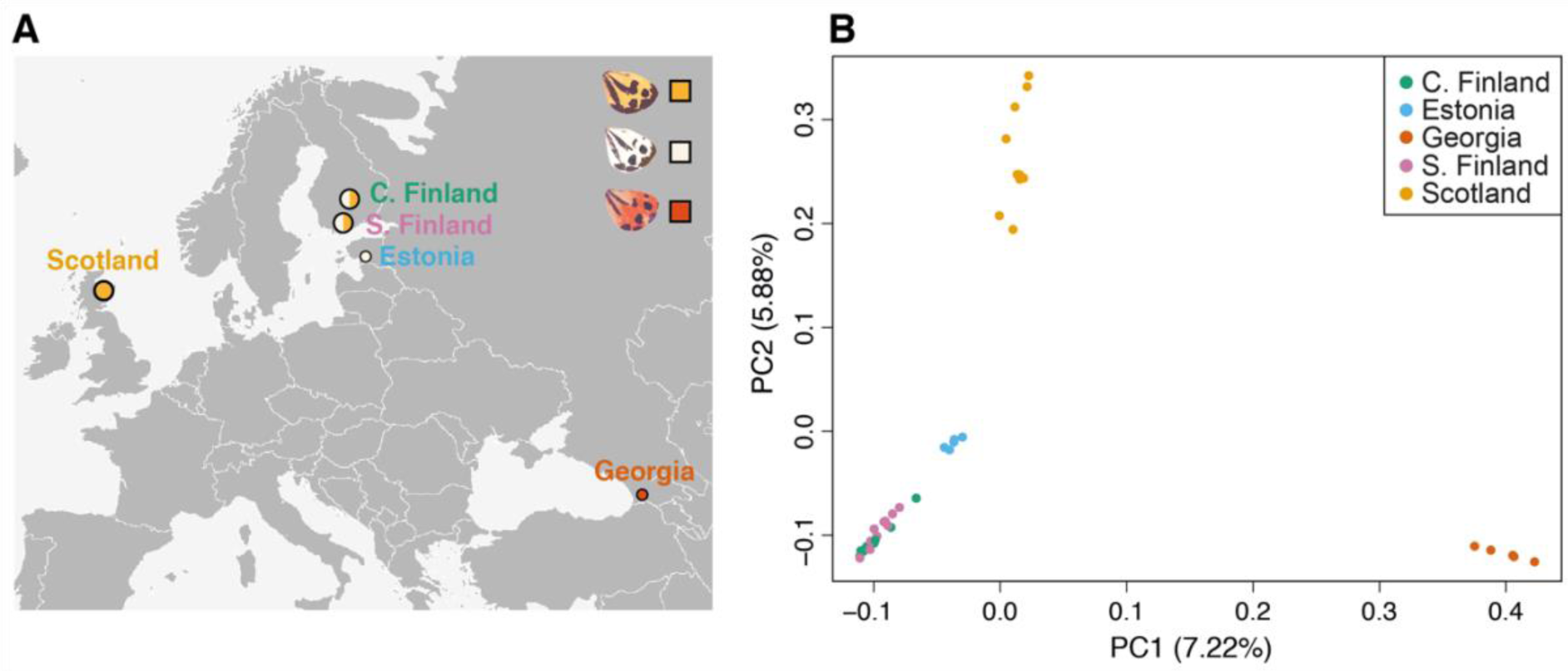
Sampling locations and population structure across *Arctia plantaginis’* European geographic range. **(A)** Sampling locations of 40 wild *A. plantaginis* males from the European portion of the Holarctic species range (see **Supplementary Table 2** for exact sampling coordinates). Circle size represents sample size (Central Finland: n=10, Estonia: n=5, Scotland: n=10, Southern Finland: n=10, Georgia: n=5), and circle colour indicates the proportion of each hindwing colour morph collected. **(B)** Genome-wide PCA (n=40; 752303 SNPs) with principle component 1 plotted against principle component 2, explaining 7.22% and 5.88% of total genetic variance, respectively.

**Figure 6.**
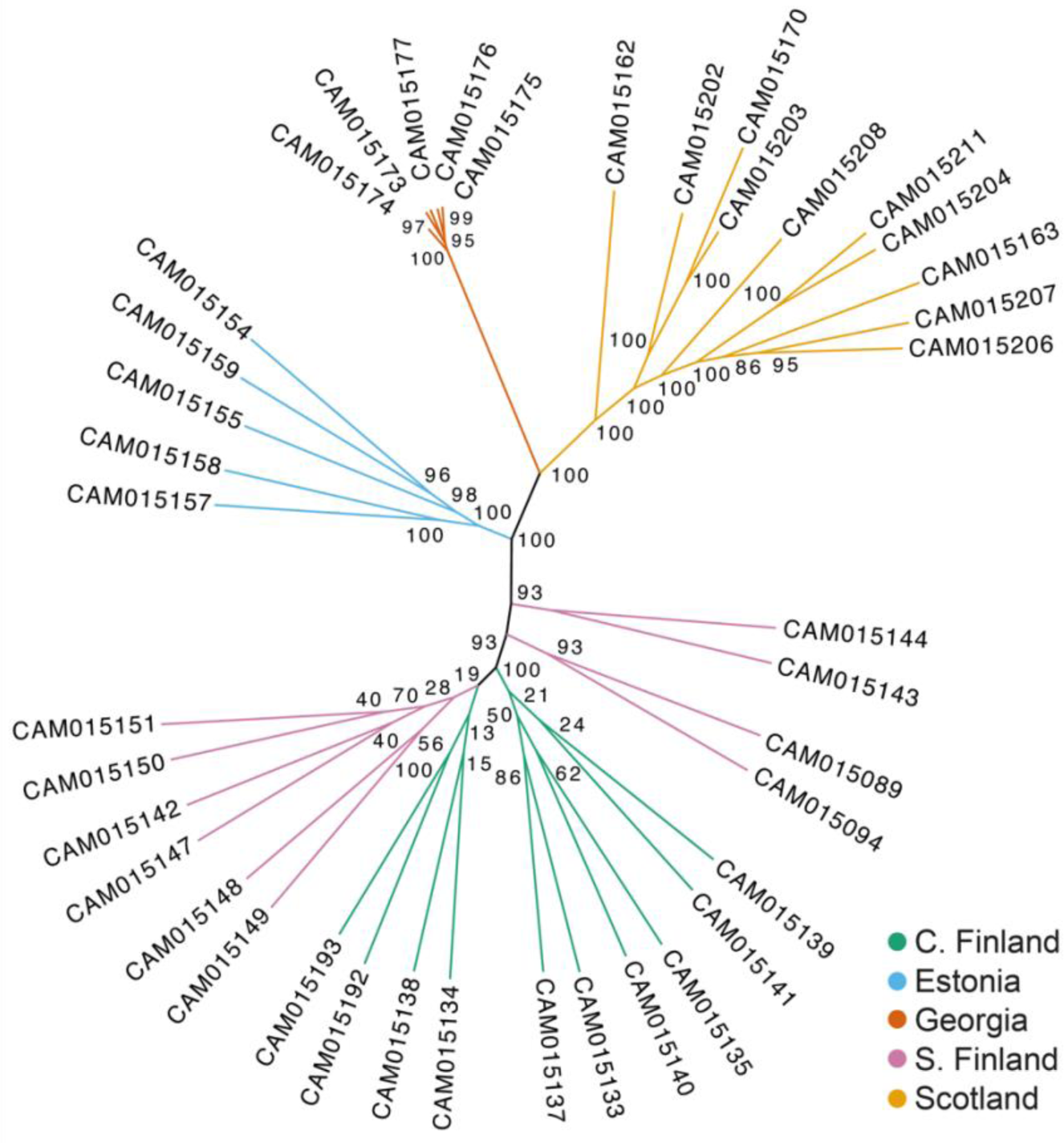
Maximum likelihood unrooted phylogeny of wild *Arctia plantaginis* males (n=40) from the European geographic range. Tree constructed using RAxML with 100 rapid bootstraps, using 558549 SNPs. Node labels indicate bootstrap support. See **Figure 5A** for sampling locations.

Internal branch lengths were strikingly shorter within the Georgian population, indicating much higher intra-population relatedness than in populations outside of Georgia (Figure 6). This signal of low genetic variation within Georgia was unlikely caused by sampling relatives, as individuals were collected from a large population. Whilst further sampling is required to confirm whether the signal persists across the Caucasus, this finding casts doubt on the hypothesis that the *A. plantaginis* species originated in the Caucasus, which is based on morphological parsimony [17]. If *A. plantaginis* spread from the Caucasus with a narrow founder population, as suggested in Hegna et al. 2015 [17], we would expect higher genetic diversity in the Caucasus compared to the other geographic regions. Similar patterns of strong genetic differentiation and low genetic diversity in Caucasus and other European mountain ranges have been observed in the Holarctic butterfly *Boloria eunomia* [72], which likely retreated into refugia provided by warmer micro-habitats within European mountain ranges during particularly harsh glaciation periods. Perhaps a similar scenario occurred in *A. plantaginis*, with founders of the Caucasus population restricted during severe glacial conditions. The species origin of *A. plantaginis* therefore remains unknown, and may be clarified by future inclusion of an *Arctia* outgroup to root the phylogenetic tree.

## Conclusions

By converting heterozygosity into an asset rather than a hindrance, trio binning provides an effective solution for *de novo* assembly of heterozygous regions, with this high-quality *A. plantaginis* reference genome paving the way for the use of trio binning to successfully assemble other highly heterozygous genomes. As the first trio binned genome available for any invertebrate species, the *A. plantaginis* assembly adds supports to trio binning as the best method for achieving fully haplotype-resolved, diploid genomes. The high-quality *A. plantaginis* reference assembly and annotation itself will contribute to Lepidoptera comparative phylogenomics by broadening taxonomic sampling into the Erebidae family, whilst facilitating genomic research on *A. plantaginis* itself.

## Availability of supporting data

All raw sequencing data for *Arctia plantaginis* reported in this article are available under ENA study accession number PRJEB36595.

## Additional files

**Supplementary Figure 1:** PacBio read length distribution for the *Arctia plantaginis* F1 offspring genome

**Supplementary Figure 2:** K-mer blob plot visualising haplotype specific k-mers for *Arctia plantaginis*

**Supplementary Figure 3**: GenomeScope profile of the *Arctia plantaginis* F1 offspring genome

**Supplementary Figure 4:** Cytogenetic analysis of *Arctia plantaginis* sex chromosomes

**Supplementary Text 1:** Cytogenetic analysis of *Arctia plantaginis* sex chromosomes

**Supplementary Table 1:** Full BUSCO summary for *Arctia plantaginis* and seven publicly available lepidopteran genome assemblies

**Supplementary Table 2:** Exact sampling localities of wild *Arctia plantaginis* males used in population genomic analysis

**Supplementary Table 3:** Resequenced genome statistics for wild *Arctia plantaginis* males used in population genomic analysis

## DECLARATIONS

## List of abbreviations

BAC: bacterial artificial chromosome,
bp: base pairs,
BUSCO: Benchmarking Universal Single-Copy Ortholog,
BWA: Burrows-Wheeler Aligner,
CLR: continuous long reads,
CTAB: hexadecyltrimethylammonium bromide,
Cy3-dUTP: cyanine 3-deoxyuridine triphosphate,
DABCO: 1,4- diazabicyclo[2.2.2]octane,
DAPI: 1,4- diazabicyclo[2.2.2]octane,
ENA: European Nucleotide Archive,
FS: Fisher strand bias,
GATK: Genome Analysis Tool Kit,
GISH: genomic *in situ* hybridization,
GTRGAMMA: generalised time-reversible substitution model and gamma model of rate heterogeneity,
KAT: Kmer Analysis Toolkit,
kbp: kilobase pairs,
LD: linkage disequilibrium,
Mbp: megabase pairs,
ml: millilitre,
ML: maximum likelihood,
MQ: root mean square mapping quality,
MQRankSum: mapping quality rank sum test,
PacBio: Pacific Biosciences,
PC: principle component,
PCA: principle component analysis,
QD: quality by depth,
RAxML: Random Axelerated Maximum Likelihood,
ReadPosRankSum: read position rank sum test,
SMRT: Single Molecule, Real-Time,
SNP: single nucleotide polymorphism,
SOR: strand odds ratio,
STAR: Spliced Transcripts Alignment to a Reference

## Ethics approval and consent to participate

Not applicable.

## Consent for publication

Not applicable.

## Author contributions

CDJ conceived and provided funding for the study. JAG and JM performed rearing and fieldwork, for which JM provided the funding. ECY and IAW performed genomic extractions. SAM performed genome assembly, for which RD provided funding. SP performed genome curation. TNG performed genome annotation. PN performed cytogenetic analysis. ECY performed comparative quality assessment. ECY performed population genomic analysis, with contributions from JIM. ECY, SAM and PN produced figures. ECY wrote the manuscript with contributions from JAG, SAM, TNG and PN, and input from all authors.

## Acknowledgements

We thank Kaisa Suisto for assisting in sample rearing. We also thank Novogene (China) for performing Illumina whole genome library preparation and sequencing, and the Wellcome Sanger Institute (Cambridge, UK) for performing PacBio and 10X Chromium library preparation and sequencing.

