## Supplementary Material for "A haplotype-resolved, *de novo* genome assembly for the wood tiger moth (*Arctia plantaginis*) through trio binning"

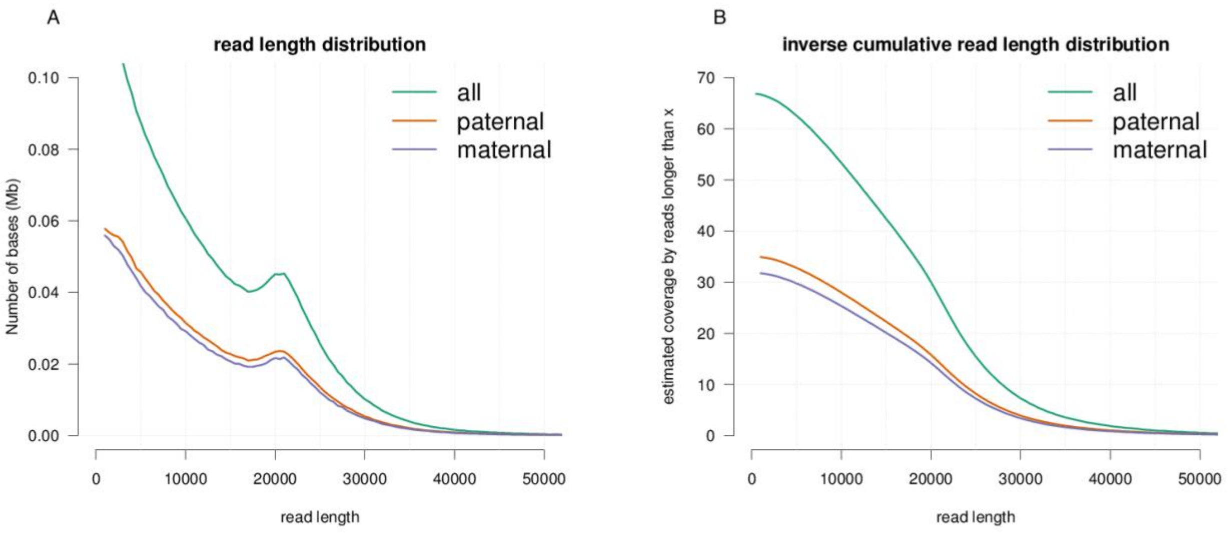


**Supplementary Figure 1**. **PacBio read length distribution for the *Arctia plantaginis* F1 offspring genome**. **(A)** Read length distribution of the entire PacBio dataset and for those reads assigned to either the maternal or paternal haplotypes. **(B)** Inverse cumulative read distribution to show coverage above a certain read length for the entire PacBio dataset and for those reads assigned to either the maternal or paternal haplotypes. A genome size of 590 Mb is assumed for the coverage estimate.

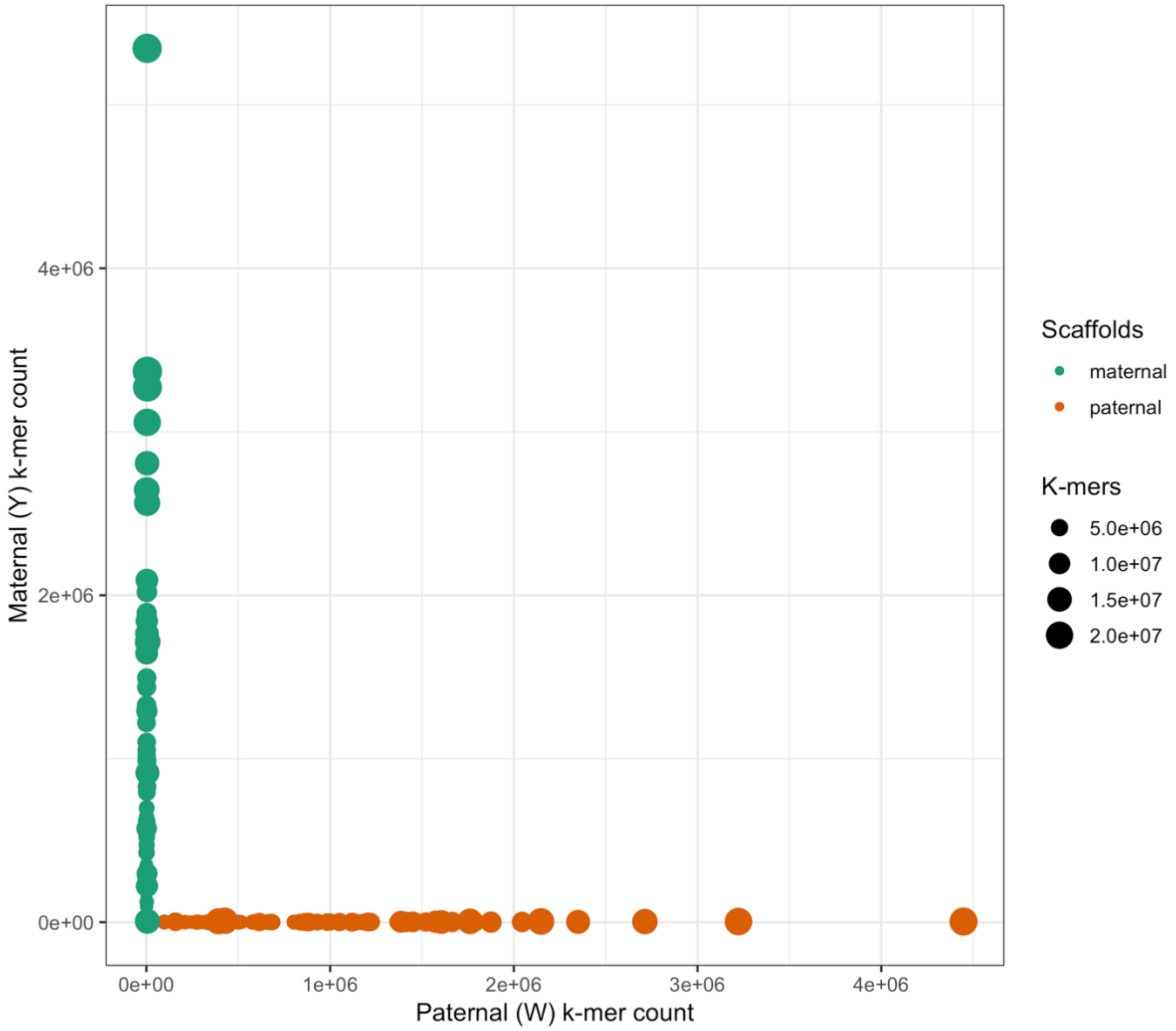


**Supplementary Figure 2. K-mer blob plot visualising haplotype specific k-mers for *Arctia plantaginis***. Plot showing for each scaffold the maternal (green) and paternal (red) assembly how many k-mers in that scaffold are found in the maternal or paternal k-mer sets. The size of the blob represents the total number of k-mers in that scaffold. There is good separation between the haplotypes, with each assembly consisting mostly of k-mers associated with the associated parental k-mer.

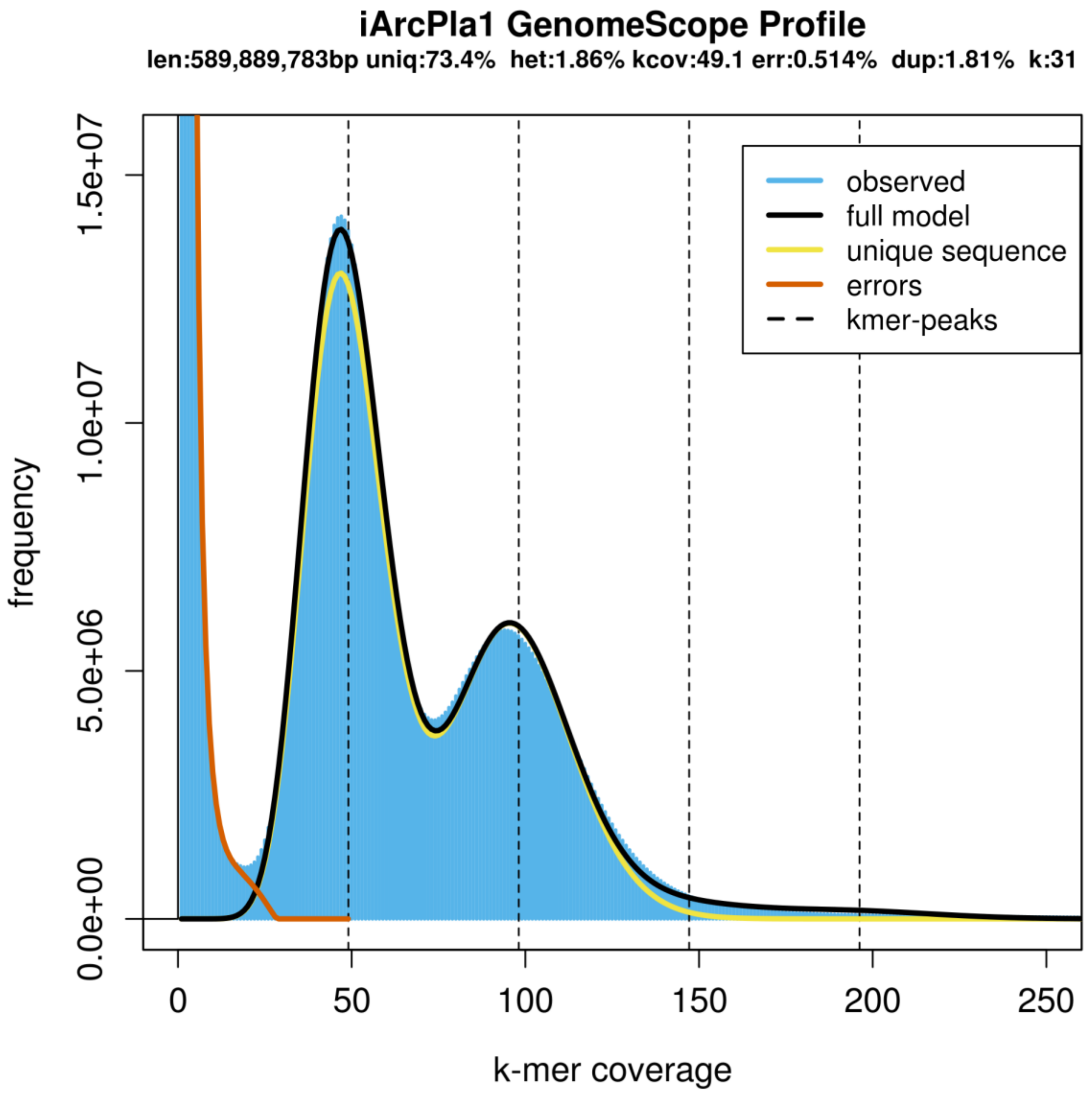


**Supplementary Figure 3. GenomeScope profile of the *Arctia plantaginis* F1 offspring genome**. K-mers are derived from 10X Genomics Illumina data. The estimated haploid genome size is 590Mb, heterozygosity 1.9% and repeat fraction 27%.

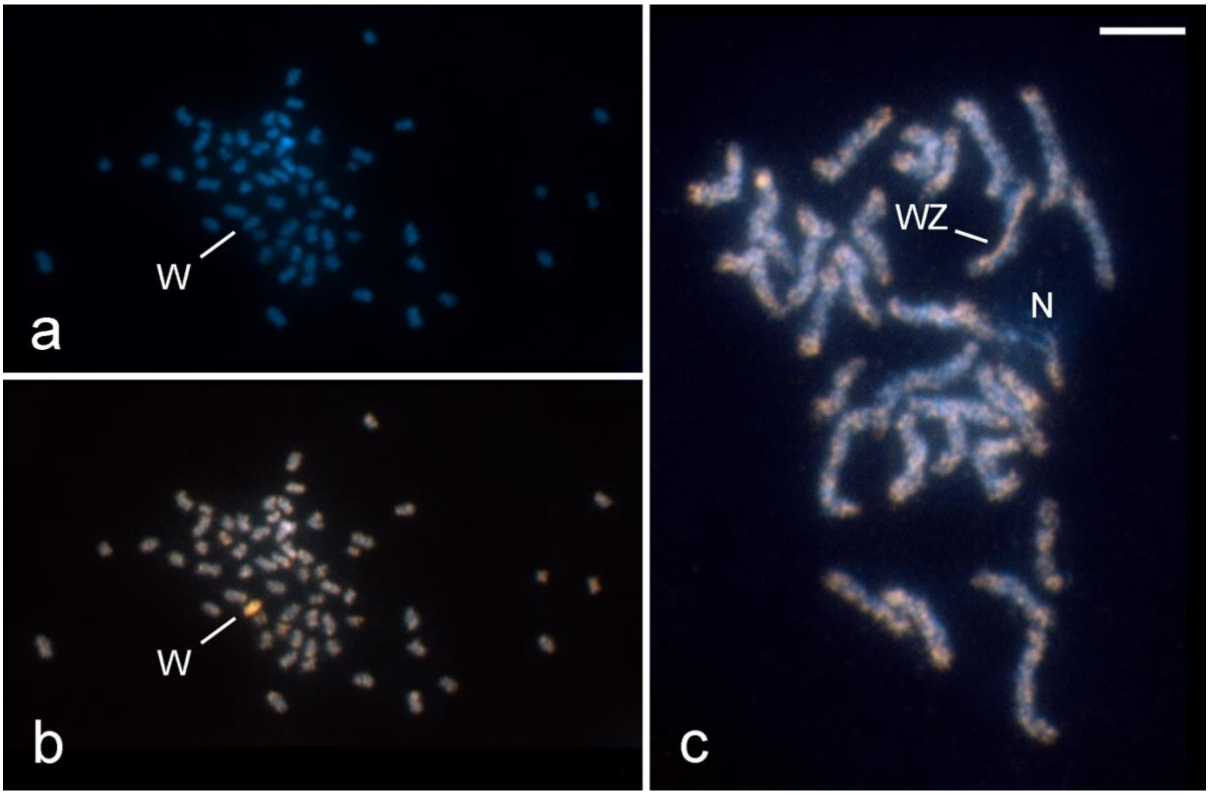


**Supplementary Figure 4. Cytogenetic analysis of *Arctia plantaginis* sex chromosomes.** Chromosomes were counterstained with DAPI (blue) and female derived genomic probe was labelled by Cy3 (orange). **(A)** Female mitotic metaphase consists of 2n=62 chromosomes. Note that the W sex chromosome, identified in **(B)**, is not highlighted by DAPI staining, indicating it is not formed by AT-rich heterochromatin. The only DAPI positive block corresponds to overlap of two chromosomes. **(B)** Genomic in situ hybridization (GISH) on the same mitotic nucleus, as in **(A)**, highlighted the W chromosome (orange). **(C)** Female pachytene nucleus consisting of n=31 bivalents were probed by GISH. The WZ sex chromosome bivalent was identified by different signal intensities of the W and Z chromosome threads. The probe (orange) painted almost the entire length of the W chromosome as well as various autosomal regions, but not the Z chromosome. This is probably due to hemizygosity of the Z chromosome in female gDNA used to construct the probe. Also note an interstitially localized nucleolus (N). Scale bar=5 µm.

**Supplementary Text 1. Cytogenetic analysis of *Arctia plantaginis* sex chromosomes.** In the female mitotic complement, DAPI staining did not highlight any chromosome which could represent the repeat-rich and thus AT-rich W chromosome (**Supplementary Figure 4A**; cf. Carabajal Paladino et al. 2019 [1]). Therefore, genomic in situ hybridization (GISH) was employed to identify the W sex chromosome in both mitotic and meiotic female nuclei (**Supplementary Figure 4B, C**). In GISH, fluorescently labelled probe derived from female gDNA is hybridized to female chromosome preparations in surplus of unlabelled male competitor gDNA. The male competitor re-associates with autosomal and Z-linked probe sequences, i.e. sequences shared by both sexes. Female specific or enriched probe sequences then preferentially hybridize to chromosomes and highlight the female specific W chromosome in both mitotic and pachytene nuclei. In *A. plantaginis*, GISH clearly identified an inconspicuous medium size chromosome as the W chromosome in female mitotic complements (**Supplementary Figure 4A, B**). The WZ sex chromosome bivalent was not identified in DAPI stained female pachytene nuclei either (not shown). This contrasts with the garden tiger moth (*Arctia caja*), in which the W chromosome was easily recognized according to an AT-rich heterochromatin block deeply stained by DAPI [2]. However, GISH with the female genomic probe labelled the W thread in the pachytene WZ sex chromosome bivalent. The hybridization signal covered almost entire W chromosome except for subtelomeric regions but also most autosomal regions (**Supplementary Figure 4C**).

| | Complete BUSCOs | Single copy BUSCOs | Duplicated BUSCOs | Fragmented BUSCOs | Missing BUSCOs |
|---|---|---|---|---|---|
| ***Arctia plantaginis*** (binned: iArcPla.TrioW, paternal haplotype) | 98.1% | 96.9% | 1.2% | 0.5% | 1.4% |
| ***Arctia plantaginis*** (binned: iArcPla.TrioY, maternal haplotype) | 96.4% | 95.3% | 1.1% | 0.5% | 3.1% |
| ***Arctia plantaginis*** (unbinned: iArcPla.wtdbg2) | 95.4% | 93.3% | 2.1% | 1.7% | 2.9% |
| ***Bicyclus anynana*** | 97.6% | 96.8% | 0.8% | 0.8% | 1.6% |
| ***Bombyx mori*** | 98.4% | 97.2% | 1.2% | 0.5% | 1.1% |
| ***Danaus plexippus*** | 98.0% | 96.0% | 2.0% | 1.0% | 1.0% |
| ***Heliconius melpomene*** | 98.7% | 97.4% | 1.3% | 0.3% | 1.0% |
| ***Manduca sexta*** | 96.7% | 93.9% | 2.8% | 2.0% | 1.3% |
| ***Melitaea cinxia*** | 83.0% | 82.9% | 0.1% | 8.5% | 8.5% |
| ***Trichoplusia ni*** | 97.4% | 96.6% | 0.8% | 1.1% | 1.5% |

**Supplementary Table 1**. **Full BUSCO summary for *Arctia plantaginis* and seven publicly available lepidopteran genome assemblies**. BUSCO analysis performed using the ‘insecta_odb9’ gene set (n=1658) to evaluate assembly completeness.

| Sample ID | Morph colour | Locality | Population | Latitude | Longitude | Altitude |
|---|---|---|---|---|---|---|
| CAM015089 | Yellow | Sandbäcken, Hanko | Southern Finland | 59.84741 | 23.132814 | 10 |
| CAM015094 | Yellow | Svanvik, Hanko | Southern Finland | 59.8393 | 23.1827 | 15 |
| CAM015133 | Yellow | Huosiaisnotko, Laukaa | Central Finland | 62.385908 | 25.818186 | 142 |
| CAM015134 | Yellow | Haralanharju, Kangasala | Central Finland | 61.534638 | 24.081058 | 99 |
| CAM015135 | Yellow | Heposuo, Laukaa | Central Finland | 62.347827 | 25.827675 | 141 |
| CAM015137 | White | Mäyrämäki, Jyväskylä | Central Finland | 62.228377 | 25.646546 | 194 |
| CAM015138 | White | Heposuo, Laukaa | Central Finland | 62.347827 | 25.827675 | 141 |
| CAM015139 | White | Lautaperä, Keuruu | Central Finland | 62.18704 | 24.874457 | 186 |
| CAM015140 | White | Heposuo, Laukaa | Central Finland | 62.347827 | 25.827675 | 141 |
| CAM015141 | White | Huosiaisnotko, Laukaa | Central Finland | 62.385908 | 25.818186 | 142 |
| CAM015142 | Yellow | Voiaskintie, Ulrikasund | Southern Finland | 60.433844 | 25.323725 | 65 |
| CAM015143 | Yellow | Tvärminne, Hanko | Southern Finland | 59.846 | 23.167 | 11 |
| CAM015144 | Yellow | Tvärminne, Hanko | Southern Finland | 59.846 | 23.167 | 11 |
| CAM015147 | White | Voiaskintie, Ulrikasund | Southern Finland | 60.433844 | 25.323725 | 65 |
| CAM015148 | White | Voiaskintie, Ulrikasund | Southern Finland | 60.433844 | 25.323725 | 65 |
| CAM015149 | White | Mosabackantie, Sipoo | Southern Finland | 60.333267 | 25.159055 | 30 |
| CAM015150 | White | Voiaskintie, Ulrikasund | Southern Finland | 60.433844 | 25.323725 | 65 |
| CAM015151 | White | Voiaskintie, Ulrikasund | Southern Finland | 60.433844 | 25.323725 | 65 |
| CAM015154 | White | Kanaküla 2 | Estonia | 58.270683 | 25.118833 | 44 |
| CAM015155 | White | Kanaküla 1.2 | Estonia | 58.263233 | 25.1449 | 45 |
| CAM015157 | White | Kanaküla 3 | Estonia | 58.301833 | 25.0969 | 36 |
| CAM015158 | White | Kanaküla 4 | Estonia | 58.2631 | 25.19075 | 56 |
| CAM015159 | White | Kanaküla 2 | Estonia | 58.270683 | 25.118833 | 44 |
| CAM015162 | Yellow | Thieves Hill, Aultmore, Keith | Scotland | 57.575232 | -3.048772 | 204 |
| CAM015163 | Yellow | Portknockie, Buckie | Scotland | 57.704217 | -2.876195 | 63 |
| CAM015170 | Yellow | Findlater Castle, Portsoy | Scotland | 57.69157 | -2.77265 | 47 |
| CAM015173 | Red | Borjomi-Kharagauli National Park | Georgia | 41.83177 | 42.83855 | 2189 |
| CAM015174 | Red | Borjomi-Kharagauli National Park | Georgia | 41.83177 | 42.83855 | 2189 |
| CAM015175 | Red | Borjomi-Kharagauli National Park | Georgia | 41.83177 | 42.83855 | 2189 |
| CAM015176 | Red | Borjomi-Kharagauli National Park | Georgia | 41.83177 | 42.83855 | 2189 |
| CAM015177 | Red | Borjomi-Kharagauli National Park | Georgia | 41.83177 | 42.83855 | 2189 |
| CAM015192 | Yellow | Heposuo, Laukaa | Central Finland | 62.347827 | 25.827675 | 141 |
| CAM015193 | Yellow | Heposuo, Laukaa | Central Finland | 62.347827 | 25.827675 | 141 |
| **CAM015202** | Yellow | Portknockie, Buckie | Scotland | 57.704217 | -2.876195 | 63 |
| **CAM015203** | Yellow | Portknockie, Buckie | Scotland | 57.704217 | -2.876195 | 63 |
| **CAM015204** | Yellow | Portknockie, Buckie | Scotland | 57.704217 | -2.876195 | 63 |
| **CAM015206** | Yellow | Portknockie, Buckie | Scotland | 57.704217 | -2.876195 | 63 |
| **CAM015207** | Yellow | Portknockie, Buckie | Scotland | 57.704217 | -2.876195 | 63 |
| **CAM015208** | Yellow | Portknockie, Buckie | Scotland | 57.704217 | -2.876195 | 63 |
| **CAM015211** | Yellow | Portknockie, Buckie | Scotland | 57.704217 | -2.876195 | 63 |

**Supplementary Table 2. Exact sampling localities of wild *Arctia plantaginis* males used in population genomic analysis**. Samples highlighted in bold are the F1 offspring of wild parents sampled from the localities presented in the table.

| Sample ID | ENA Sample Accession Number | Mean sequencing coverage | Number of raw reads | % of raw reads mapped against iArcPla.TrioW |
|---|---|---|---|---|
| CAM015089 | ERS4285276 | 13.3X | 67913274 | 99.02% |
| CAM015094 | ERS4285277 | 13.6X | 71591177 | 98.88% |
| CAM015133 | ERS4285280 | 16.0X | 91510105 | 98.94% |
| CAM015134 | ERS4285281 | 14.9X | 84005409 | 99.03% |
| CAM015135 | ERS4285282 | 14.4X | 82119542 | 99.00% |
| CAM015137 | ERS4285283 | 16.0X | 90270209 | 98.99% |
| CAM015138 | ERS4285284 | 14.7X | 83204225 | 98.98% |
| CAM015139 | ERS4285285 | 15.9X | 91852724 | 99.04% |
| CAM015140 | ERS4285286 | 15.1X | 85825617 | 98.96% |
| CAM015141 | ERS4285287 | 13.5X | 76558659 | 99.01% |
| CAM015142 | ERS4285288 | 15.0X | 84287184 | 99.04% |
| CAM015143 | ERS4285289 | 14.4X | 81283183 | 99.00% |
| CAM015144 | ERS4285290 | 15.3X | 88352550 | 98.97% |
| CAM015147 | ERS4285291 | 13.6X | 75042444 | 99.00% |
| CAM015148 | ERS4285292 | 16.0X | 89777636 | 99.00% |
| CAM015149 | ERS4285293 | 13.2X | 73604268 | 99.03% |
| CAM015150 | ERS4285294 | 13.0X | 72332606 | 98.98% |
| CAM015151 | ERS4285295 | 13.9X | 80159712 | 99.06% |
| CAM015154 | ERS4285296 | 12.9X | 71746939 | 98.93% |
| CAM015155 | ERS4285297 | 15.6X | 90247219 | 98.99% |
| CAM015157 | ERS4285298 | 14.4X | 82512306 | 98.99% |
| CAM015158 | ERS4285299 | 13.4X | 77179045 | 98.98% |
| CAM015159 | ERS4285300 | 13.9X | 79287964 | 98.99% |
| CAM015162 | ERS4285301 | 14.5X | 82886426 | 98.95% |
| CAM015163 | ERS4285302 | 14.7X | 83217243 | 98.94% |
| CAM015170 | ERS4285303 | 10.6X | 59511526 | 98.92% |
| CAM015173 | ERS4285304 | 11.1X | 68516047 | 98.09% |
| CAM015174 | ERS4285305 | 10.5X | 65751644 | 98.50% |
| CAM015175 | ERS4285306 | 12.1X | 75845780 | 98.45% |
| CAM015176 | ERS4285307 | 12.1X | 74743346 | 98.46% |
| CAM015177 | ERS4285308 | 13.2X | 83009022 | 98.44% |
| CAM015192 | ERS4285309 | 14.0X | 80351585 | 98.85% |
| CAM015193 | ERS4285310 | 10.6X | 58462162 | 99.04% |
| CAM015202 | ERS4285311 | 11.0X | 59903009 | 99.00% |
| CAM015203 | ERS4285312 | 10.1X | 57251863 | 99.03% |
| CAM015204 | ERS4285313 | 11.6X | 64780636 | 98.08% |
| CAM015206 | ERS4285314 | 15.0X | 87719259 | 98.90% |
| CAM015207 | ERS4285315 | 11.1X | 62197559 | 96.13% |
| CAM015208 | ERS4285316 | 11.9X | 64848486 | 98.96% |
| CAM015211 | ERS4285317 | 12.0X | 63799250 | 99.04% |

**Supplementary Table 3. Resequenced genome statistics for wild *Arctia plantaginis* males used in population genomic analysis.**

### Supplementary Material References

1. Carabajal Paladino LZ, Provazníková I, Berger M, et al. Sex Chromosome Turnover in Moths of the Diverse Superfamily Gelechioidea. Genome Biol Evol. 2019; 11: 1307–1319.

2. Nguyen P, Sahara K, Yoshido A, Marec F. Evolutionary dynamics of rDNA clusters on chromosomes of moths and butterflies (Lepidoptera). Genetica. 2010; 138: 343–354.
